# The second edge of ADHD: an advantage in motor learning and performance with task-irrelevant background vibratory noise

**DOI:** 10.1101/426916

**Authors:** Maria Korman, Lian Meir-Yalon, Nebal Egbarieh, Avi Karni

**Affiliations:** The Edmond J. Safra Brain Research Center for the Study of Learning Disabilities, University of Haifa, Haifa, Israel; Laboratory for Human Brain and Learning, Sagol Dept. of Neurobiology, University of Haifa, Haifa, Israel; Department of Occupational Therapy, Faculty of Social Welfare & Health Sciences, University of Haifa, Israel

**Author notes:** Equal contribution (Maria Korman, Lian Meir-Yalon). **Conflict of interest:** The authors declare no competing financial interests. Corresponding author: Maria Korman University of Haifa 199 Aba Khoushy Ave. Mount Carmel, Haifa ISRAEL.

**Keywords:** procedural learning, motor sequence, skill memory consolidation, ADHD, chronotype, arousal, young adults, sensory stimulation

## Abstract

Young adults with ADHD often gain less than expected from practice sessions well-suited for their peers. Here, we tested whether task-irrelevant, low-intensity vibratory stimulation (VtSt), suggested to modulate motor learning, may compensate for such learning deficits. Participants were given training, either with or without VtSt, on a sequence of finger opposition movements. Under VtSt, typical individuals had reduced overnight, consolidation phase, gains; performance partly recovering one week later. In contrast, participants with ADHD benefitted from VtSt both during the acquisition (online) and the overnight skill consolidation (offline) phases. One week later, both groups showed robust retention of the gains in performance, but when tested with background VtSt, individuals with ADHD outperformed their typical peers. We propose that ADHD can confer advantages in performance, learning and skill memory consolidation in specific ‘noisy’ conditions that adversely affect typical adults; we conjecture that the effects of VtSt are contingent on baseline arousal levels.

---

The evidence from skill acquisition studies in people with ADHD is equivocal; some studies report deficits vis-à-vis typical controls ^1-3^ while in other tasks, participants with ADHD were as effective learners as their typical peers^4,5^. Repeated task performance is essential for the acquisition of daily and academic skills, but, long repetitive practice can be sub-optimal in ADHD^2,6,7^, presumably due to difficulties in sustaining attention ^8^. Deficits in directing, focusing and maintaining attention^9^ may account for the increases in error rates in ADHD^1-3^. Nevertheless, the learning of an implicit movement sequence (SRT task) in adults with ADHD was found intact^3^. In explicit learning conditions, both the acquisition and the memory consolidation phase after motor practice may be atypical in individuals with ADHD (smaller and/or slower when compared to controls), however, clear practice related gains and effective retention of the acquired skills were reported^1,10^.

Brain plasticity, the basis for skill and knowledge, is a highly controlled (selective) process, mainly because of a consolidation phase, wherein structural modifications occur at brain areas engaged in task performance and in circuits wherein the memory was initially encoded during salient experiences^11^. In the context of skill (procedural, ‘how to’) learning, these processes are triggered by the learning experience, if sufficient practice is afforded^12^. Once triggered, consolidation processes can proceed ‘off-line’, during both wakefulness and sleep, and culminate in the establishment of new knowledge and its integration into previously existing knowledge^11,13-15^. This is reflected in behavior. Large gains in performance speed, with no loss of accuracy, occur early in training, within session (‘fast learning’, novelty, phase) ^15-17^. However, additional robust gains in speed and accuracy can be expressed hours after the termination of training, for example by 24 hours post-training. These delayed (between-sessions, ‘offline’) gains in performance presumably reflect the latent neuronal long-term memory consolidation processes^14,15,17-19^. The performance level attained after the completion of the consolidation phase can be well retained for weeks and months^15^. However, the triggering or completion of a consolidation phase may fail; for example, in cases when practice is terminated too early^13,20^ when interference by subsequent experiences takes place^15,21^ and by poor sleep^22^.

Most models of memory, at the level of brain mechanisms, focus on the neural events directly (in a causal sense) mediating memory, e.g., synaptic consolidation, and the anatomical locus of the ‘memory trace’ in relation to the learning experience^11,23^. There is, however, evidence indicating that the generation of long-term memory is modulated and controlled by factors that relate to the background brain states during and after the learning experience rather than to parameters intrinsic to the training experience per se^24^. Thus, processes that are in a sense orthogonal to the actual learning experience can nevertheless gate and determine long-term memory storage^25^. Within training and post-training treatments, pharmacological and behavioral, were shown to selectively enhance or impair memory storage in many learning tasks^15,26-28^. For example, minor vibrotactile or vibroauditory stimulation afforded during training may disrupt consolidation processes in healthy young adults^29^. Thus, the actual learning experience, while obligatory, may not by itself suffice for establishing long-term memory; control mechanisms must be satisfied before learning can be consolidated into long-term memory.

Recently, a number of non-pharmacological interventions to up-regulate skill learning in ADHD were suggested. One line of evidence suggests that some of the relative learning deficits in persons with ADHD, could be corrected when training was shortened^1,2,6,30^ presumably decreasing the burden of long repetitive practice on mechanisms of sustained attention^8^. An additional line of evidence indicates that motor training scheduled to evening hours can enhance off-line memory consolidation (the expression of delayed learning gains) in young adults with ADHD and close their learning gaps vis-à-vis typically developing adults^10^; presumably because evening hours are the optimum performance hours for evening-type individuals, a chronotype characterizing many of the individuals with ADHD^31^. It was also shown that five minutes of vigorous physical activity improve affect and executive functioning of children with symptoms of ADHD^32^.

The beneficial effects of both time-of-day and physical activity may reflect an effect of the general level of arousal during practice in the acquisition of skill in persons with ADHD. Theoretical accounts of ADHD, such as the state regulation model^33^ and dual-process models^34,35^ propose that the high within-subject fluctuations of cognitive performance in ADHD may reflect problems in regulating arousal^36,37^. An optimal arousal level is considered a prerequisite for successful cognitive functioning, as both too little or too much arousal can adversely affect task performance^38,39^. Individuals with ADHD tend to be under-aroused in “normal” performance^40,41^ and learning conditions^39,42,43^.

Arousal levels are affected by environmental noise^44^. Task-irrelevant sensory noise is ubiquitous and is mostly considered detrimental and distractive^29,45^; individuals with ADHD can be even more prone to distraction than typical peers^46^. Nevertheless, improvements in the performance of individuals with ADHD were reported in various primary tasks when extra-task stimulation, such as auditory noise, was added (e.g.,^47-50^). These paradoxical effects are not well understood, but background sensory stimulation was suggested to serve as a generator of increased arousal^39^ or as a compensatory input needed to upregulate a hypo-functioning dopaminergic system in ADHD^51,52^.

The objective of the current study was to compare the immediate and long-term effects of low-intensity, task-irrelevant ‘noise’ - vibro-tactile stimulation to the trunk combined with acoustic vibration through earphones - afforded during the practice of an instructed finger opposition sequence (FOS) in non-medicated young adults with ADHD and their typical peers (without ADHD) (Figure 1). The FOS task was used as the to-be-learned task because numerous studies have shown that the time-course of FOS learning in young adults with ADHD is atypical^1,10,30,53^. Behavioural measures of speed and accuracy of performance at successive time points following a single training session (immediate, 24h and one week re-tests) were assessed. We tested the conjecture that the background vibratory stimulation (VtSt) would act as a non-specific stimulant for the ADHD group and would therefore enhance motor performance both during the acquisition and the consolidation phases. In typical young adults, with no ADHD, VtSt was recently found to adversely affect the consolidation gains in FOS performance^10^. We also tested performance with or without VtSt afforded during re-testing at one week post-training; the conjecture was that if VtSt would become, at least in part, an integrated aspect in the skill attained in practice, VtSt affordance would significantly upregulate performance.

**Figure 1:**
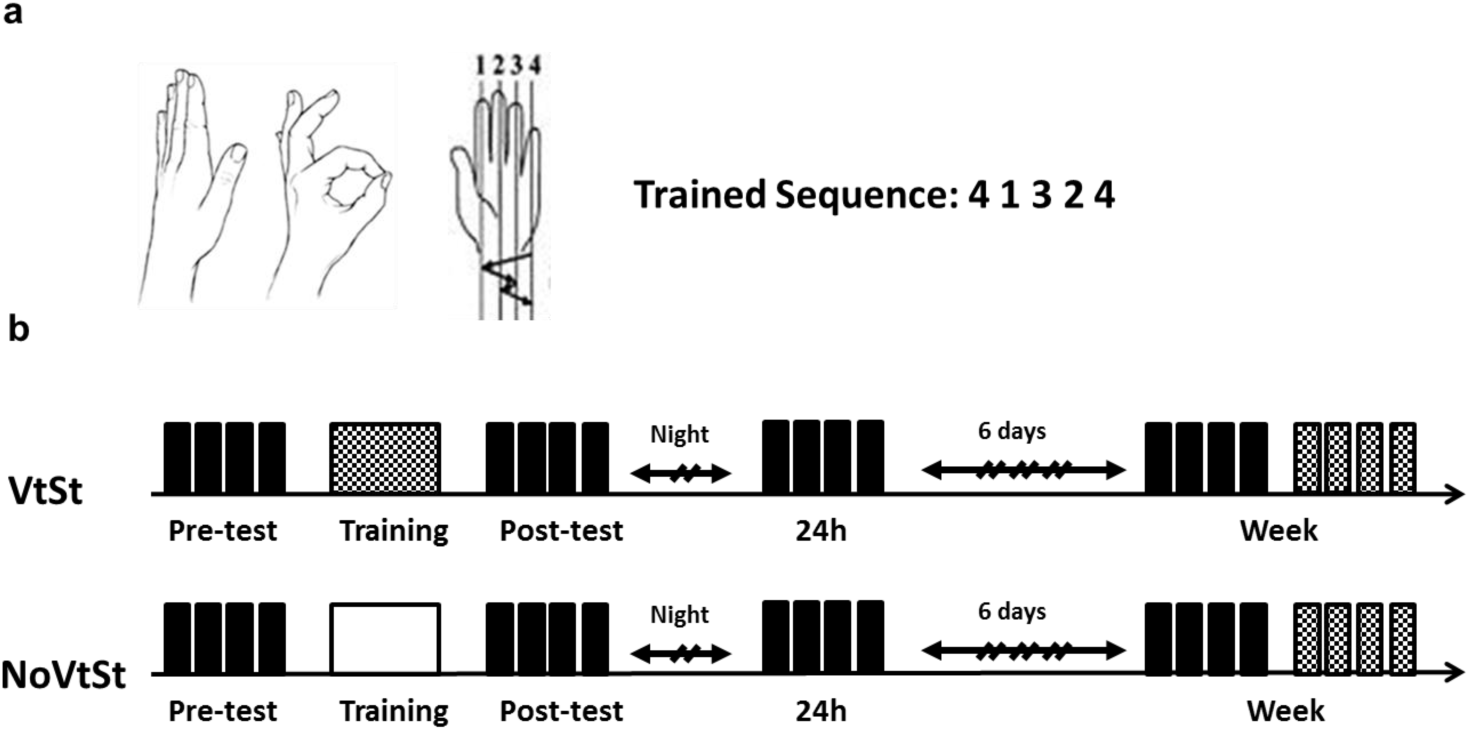
Study task and design. a) The finger-to-thumb opposition sequence (FOS) task. b) Participants were trained (160 cued repetitions of the sequence) without VtSt (white box) or with VtSt (checkboard box) in a single practice session in the morning hours. Performance was tested (four 30-sec. self-initiated performance blocks without VtSt) in 4 time-points: Pre-test before training, Post-test immediately after training, 24h after training and a Week after training (black narrow boxes). Performance of the trained sequence was also re-tested with VtSt afforded concurrently with the test blocks (checkboard narrow boxes).

Participants from both pools, ADHD and Control, were randomly assigned to one of the two experimental conditions: (1) Training with vibrotactile sensory stimulation: ADHD group, N=17 (ADHDVtSt) and Control group, N=16 (ContVtSt); (2) Training without vibrotactile sensory stimulation (ADHDNoVtSt, N=16 and ContNoVtSt, N=16) (Figure 1b). During the training blocks the participants in the sensory stimulation groups (VtSt) experienced minor vibrations delivered to the trunk by means of a commercial vibrating cushion (Homedics Inc). The cushion produced vibrotactile stimulation with a main frequency of ~65Hz, resulting in ~41dB noise.

## Results

### Absolute data

There were no significant differences in performance speed (the number of correct sequences performed, on average, in the test) between the four groups (one-way ANOVA, F(3,64)=0.478, p=0.699) at pre-training (pre-test). Also, there was no significant difference between the pre-training performance of the participants with and without ADHD (t(63)=-0.668; p=0.507).

Overall, there was a significant improvement in speed across the study period (4 time-points), in all four groups (F(3,183)=330.351, p<0.001, MSE=2930.21, η^2^=0.844) (**Figure 2a**). There was no significant group effect (p=0.465), but there was a trend towards a significant interaction of time-point X group (F(9,183)=2.108, p=0.069, MSE=16.579, η^2^=0.082). Post-hoc group comparisons showed that participants with ADHD when trained without background stimulation (ADHDNoVtSt) gained relatively less, overall, compared to the ADHDVtSt and the ContNoVtSt groups, although their performance did not significantly differ from that of typical young adults experiencing the background stimulation (ContVtSt) **(Figure 2a,b).**

**Figure 2.**
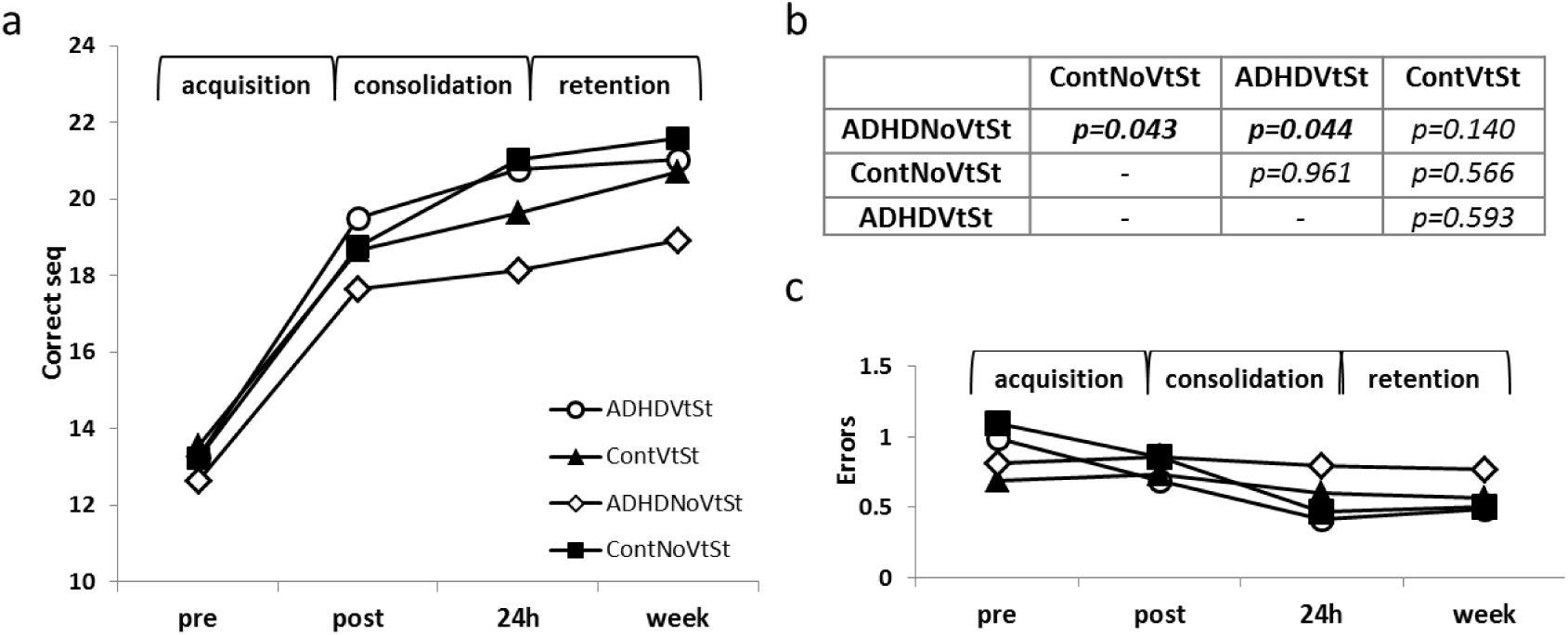
The time course of performance changes in the four experimental groups (ADHDNoVtSt-ADHD training with no vibratory sensory stimulation, ADHDVtSt - ADHD training with vibratory sensory stimulation, ContNoVtSt - typical adults training with no vibratory sensory stimulation, ContVtSt - typical adults training with no vibratory sensory stimulation). (**a**) The number of correct sequences tapped in a test block (speed) in the 4 time-points. All groups benefitted from training and improved across the study period. The acquisition and “offline”, consolidation phase gains in speed were well maintained across the 1 week retention interval. (**b**) Post-hoc (LSD) pair-wise group comparisons of performance speed across the four time points. Participants with ADHD given training without background stimulation (ADHDNoVtSt) differed from the ADHDVtSt and the ContNoVtSt groups in the time-course of improvement. (**c**) The absolute number of sequencing errors committed in the 4 tests (time-points). Performance was very accurate throughout. Each data point depicts the mean of group performance at the time-point; bars - SEM.

Tests to assess the contributions of the 3 time intervals (acquisition; overnight consolidation and 1-week retention) to the improvements in speed, showed that the training session resulted in early (within-session) gains and in additional delayed (post-training, time-dependent) gains in performance, across all groups (**Figure 2a**). The within-session gains were robust and similar across groups *(acquisition interval*: F(1,61)=443.106, p<0.001, MSE=3579.40, η^2^=0.879) and there was no significant group x time-point interaction (F(1,61)=1.252, p=0.299, MSE=10.117, η^2^=0.058). The delayed, off-line, gains in performance were robust as well *(consolidation interval:* F(1,61)=36.317, p<0.001, MSE=252.972, η^2^=0.373). However, there was a significant group x time-point interaction during this phase (F(3,61)=3.221, p=0.029, MSE=22.439, η^2^=0.137), reflecting a relative lag that developed by 24h post-training in the ADHDNoVtSt group compared to the control participants trained with or without the background stimulation; this lag was apparent also relative to the participants with ADHD who were afforded VtSt during training (**Figure 2**). The gains in speed attained at the 24h post-training test were well maintained over the 1-week retention interval with small but significant further improvements *(retention interval:* F(1,61)=10.374, p=0.002, MSE=47.624, η^2^=0.145). No significant group effects (p=0.159) or a group X time-point interaction (F(1,61)=1.102, p=0.355, MSE=5.059, η^2^=0.051)were observed.

On average the participants in all four groups tended to commit very few, if any, errors (**Figure 2c**). Nevertheless, absolute accuracy improved significantly across the study period (F(3,183)=6.256, p<0.001, MSE=6.527, η^2^=0.093) i.e., the absolute number of errors decreased in all 4 groups. Thus, there was no trade-off between the improvements in speed and the number of errors committed. There was no significant group effect (p=0.141) and no significant interaction of time-point X group (p=0.251).

The effects of VtSt, afforded during training, were further explored in participants with ADHD. There were no significant differences in the initial performance (pre-test) of the two ADHD groups (t(31)=-0.633, p=0.532 and t(30)=-0.658, p=0.515; speed and accuracy, respectively), and moreover both groups (ADHDNoVtSt, ADHDVtSt) showed, across the 4 time-points, significant gains in speed (correct sequences) and a decrease in errors (F(3,93)=141.318, p<0.001, MSE=1379.324, η^2^=0.820; F(3,93)=2.651, p=0.053, MSE=2.516, η^2^=0.079, respectively) (**Figure 3a**). There was a trend towards a significant main effect of group for correct sequences (F(1,31)=3.347, p=0.077, MSE=411.658, η^2^=0.0.97) but not for errors (p= 0.303). There was also a trend towards a significant time-point X group interaction in speed (F(3,93)=2.428, p=0.070; MSE=23.695, η^2^=0.073) but not for errors (p=0.114).

**Figure 3.**
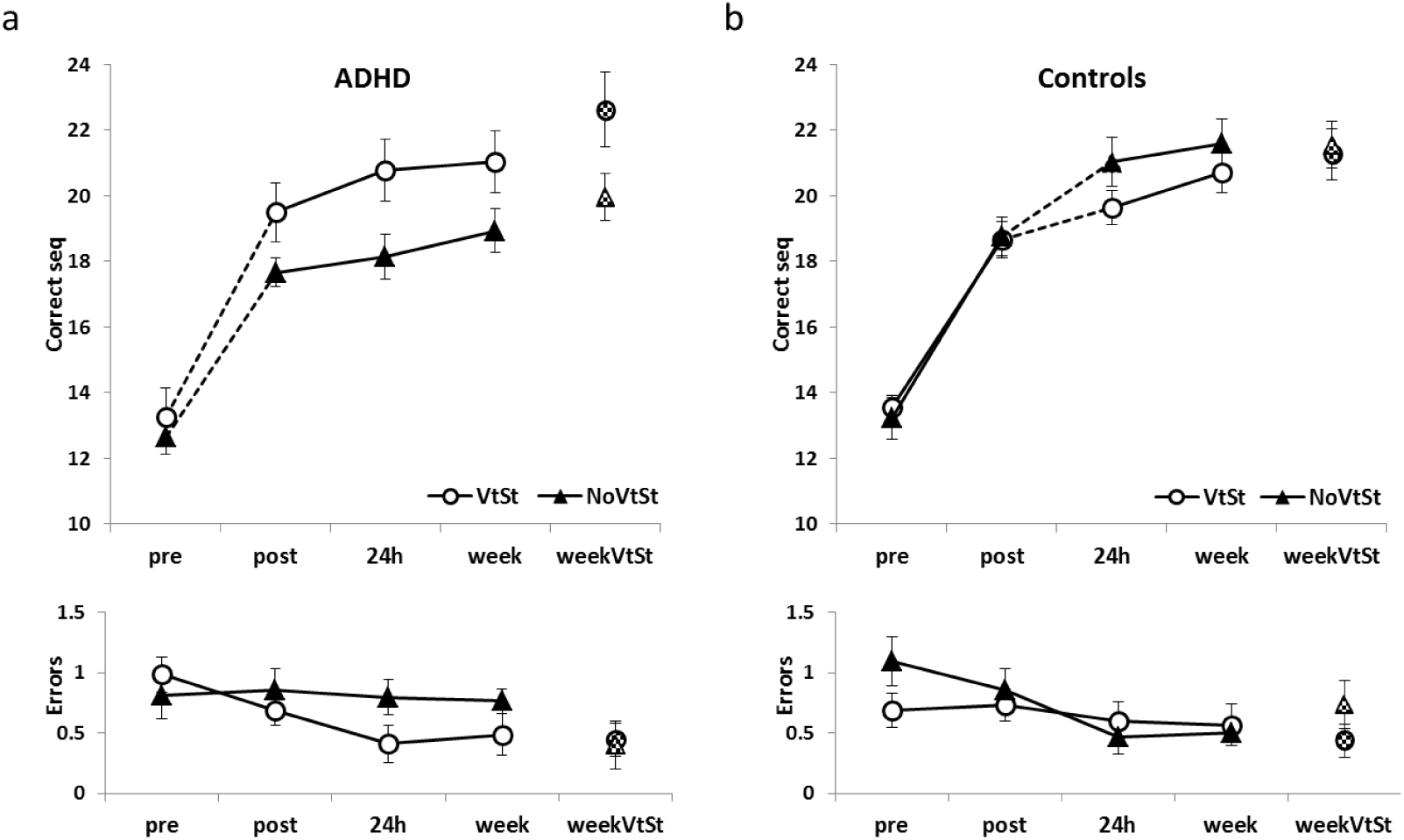
Time-course of learning with (open circles) or without (black triangles) background sensory stimulation. (**a**) Participants with ADHD. Training with VtSt resulted in larger within-session gains in the number of correct sequences and this advantage of the ADHDVtSt group was further increased by robust between-session gains attained at the 24h post-test. Both ADHD groups showed a robust boost in performance when VtSt was afforded during the retention test. (**b**) Control participants. There were robust within-session gains in the number of correct sequences, irrespective of whether VtSt was afforded, but training with VtSt resulted in smaller between-session gains. Both Control groups were unaffected by VtSt afforded during the retention test. Upper panels - mean number of correct sequences; Lower panels - number of errors. Dashed line - significant groups X time-point interaction; Chess board markers - test with VtSt; bars - SE.

The two ADHD groups expressed similar gains across the training session. Both groups improved in speed, with no costs in accuracy (speed: F(1,31)=243.870, p<0.001, MSE=2021.758, η^2^=0.887; accuracy: p=0.259) and there were no significant group effects (p=0.195; p=0.992, speed and accuracy, respectively). There was a trend towards a significant group X time-points interaction for correct sequences, (F(1,31)=3.323, p=0.078, MSE=26.329, η^2^= 0.097) (but not for errors, p=0.127) reflecting the larger gains in performance rates in the ADHDVtSt compared to the ADHDNoVtSt group (**Figure 3a**). One-way ANOVA confirmed that at immediate post-training test there was a marginally significant advantage in performance rate for the participants with of the VtStADHD group (speed: F(1,31)=3.391, p=0.075, MSE=27.381; accuracy: p=0.650).

Participants with ADHD continued to improve across the 24h between-sessions interval (comparing post-test, 24h). There were robust additional improvements in terms of speed with no loss of accuracy (F(1,31)=10.047, p=0.003, MSE=70.375, η^2^= 0.245; p=0.129, respectively). Although no significant group X time-point interactions were found (p=0.220, p=0.330, speed and accuracy, respectively), there was a significant group effect (F(1,31)=4.740, p=0.037, MSE=328.115, η^2^=0.133). Indeed, one-way ANOVA confirmed that at 24h post-training there was a significant advantage in performance rate for the participants with ADHD given the VtSt during training (speed: F(1,31)=5.226, p=0.029, MSE=57.394; accuracy: p=0.123).

There was a trend for additional gains across the retention interval (comparing 24h test, 1-week test) in speed (F(1,31)=2.922, p=0.097, MSE=17.531, η^2^=0.086) with no loss of accuracy (p=0.851). There was no significant group X time-point interaction for speed or accuracy (p=0.237, p=0.642, respectively), but the advantage of the ADHDVtSt group persisted. It was reflected in significant group effects both for the number of correct sequences executed (F(1,31)=4.572, p=0.040, MSE=371.364, η^2^=0.129) and the number of errors (F(1, 31)= 5.370, p=0.027, MSE=7.300, η^2^=0.148), with the ADHDVtSt group outperforming the ADHDNoVtSt group (**Figure 3a**).

The effects of adding the background sensory stimulation (VtSt), on the performance of the trained movement sequence, were assessed during the 1-week retention test (**Figure 3a**). The affordance of VtSt during the performance test significantly improved the performance both in terms of the number of correct sequences (increased) and of the number of errors committed (decreased) (speed: F(1,31)=15.995, p<0.001, MSE=107.088, η^2^=0.348 and accuracy: F(1,31)=6.017, p=0.020, MSE=3.116, η^2^=0.163). A trend towards a significant group effect was also found, only for speed (F(1,31)=3.588, p=0.068, MSE=354.708, η^2^=0.107; accuracy: p=0.436). There was no significant group X time-point interaction for speed (p=0.424); accuracy tended to improve in the ADHDNoVtSt group (F(1,31)= 3.823, p=0.060, MSE=1.979, η^2^=0.110).

Similar analyses in participants without ADHD (Control groups), showed that the affordance of VtSt during training had no effect on the immediate post training performance, but resulted in relatively smaller gains in speed and accuracy expressed during the overnight, 24 hours consolidation phase; in line with a previous study^29^. Nevertheless, the gap between the two groups tended to close by 1 week post training (**Figure 3b**). There were no significant differences in initial performance (pre-test) between the two Control groups (t(30)=-0.451, p=0.655, speed; t(30)=1.650; p=0.109, accuracy) and both groups (ContNoVtSt and ContVtSt) showed significant gains (F(3,90)=196.252, p<0.001, MSE=1560.55, η^2^=0.867 and F(3,90)=3.636, p=0.016, MSE=4.148, η^2^=0.053, speed and accuracy, respectively) across the four time-points of the study. There was no main effect of group for correct sequences (p=0.505) and for errors (p= 0.637), but there was a marginally significant time-point X group interaction in speed (F F(3,90)=2.379, p=0.075, MSE=18.913, η^2^=0.073) though not for accuracy (p=0.176). A comparison of the performance of the two Control groups for each of the study phases (acquisition, 24 hours consolidation, 1-week retention) showed robust learning and no differences between groups during acquisition *(Supplementary Results)* (**Figure 3b**). However, the ContVtSt group had smaller consolidation phase gains compared to the ContNoVtSt group. Both control groups continued to improve across the 24h between-sessions interval in terms of speed (F(1,30)=22.893, p<0.001, MSE=167.379, η^2^=0.433) and at no cost in accuracy (f(1,30)=3.839, p=0.059, MSE=4.254, η^2^=0.113). But although no significant group effects were found (p=0.360, p=0.892; speed and accuracy, respectively) there was a trend towards a group X time-point interaction for speed (F(1,30)=3.681, p=0.065, MSE=26.910, η^2^2= 0.109), reflecting the smaller, on average, consolidation phase gains expressed when VtSt was afforded during training. There was no interaction effect for the errors (p=0.380). In both control groups, additional gains in speed occurred during the week-long retention interval (F(1,30)=14.325, p=0.001, MSE=42.250, η^2^=0.323) with no costs in accuracy (p=0.999).

The affordance of VtSt during the performance test did not affect the performance of control subjects both in terms of the number of correct sequences (p=0.431) and of the errors (p=0.755). There were no group effects (p=0.554 and p=0.510; speed and accuracy, respectively) and no significant group X time-point interaction (p=0.378 and p=0.218, speed and accuracy, respectively).

### Normalized data

To enable a direct comparison between the gains of the ADHD and the no-ADHD groups, data were normalized relative to the mean pre-test baseline performance of each individual, yielding the relative improvements of each individual for the acquisition, the overnight consolidation and the retention intervals **(Figure 4)**. There were no significant differences in acquisition phase gains between the NoVtSt groups (ContNoVtSt, ADHDNoVtSt; (t(30)=-0.156, p=0.649). However, the acquisition gains in the 2 groups experiencing VtSt during training tended towards a significant difference (t(31)=-1.796, p=0.080), with the ADHDVtSt group tending on average to outperforming the ContVtSt group **(Figure 4a)**.

**Figure 4.**
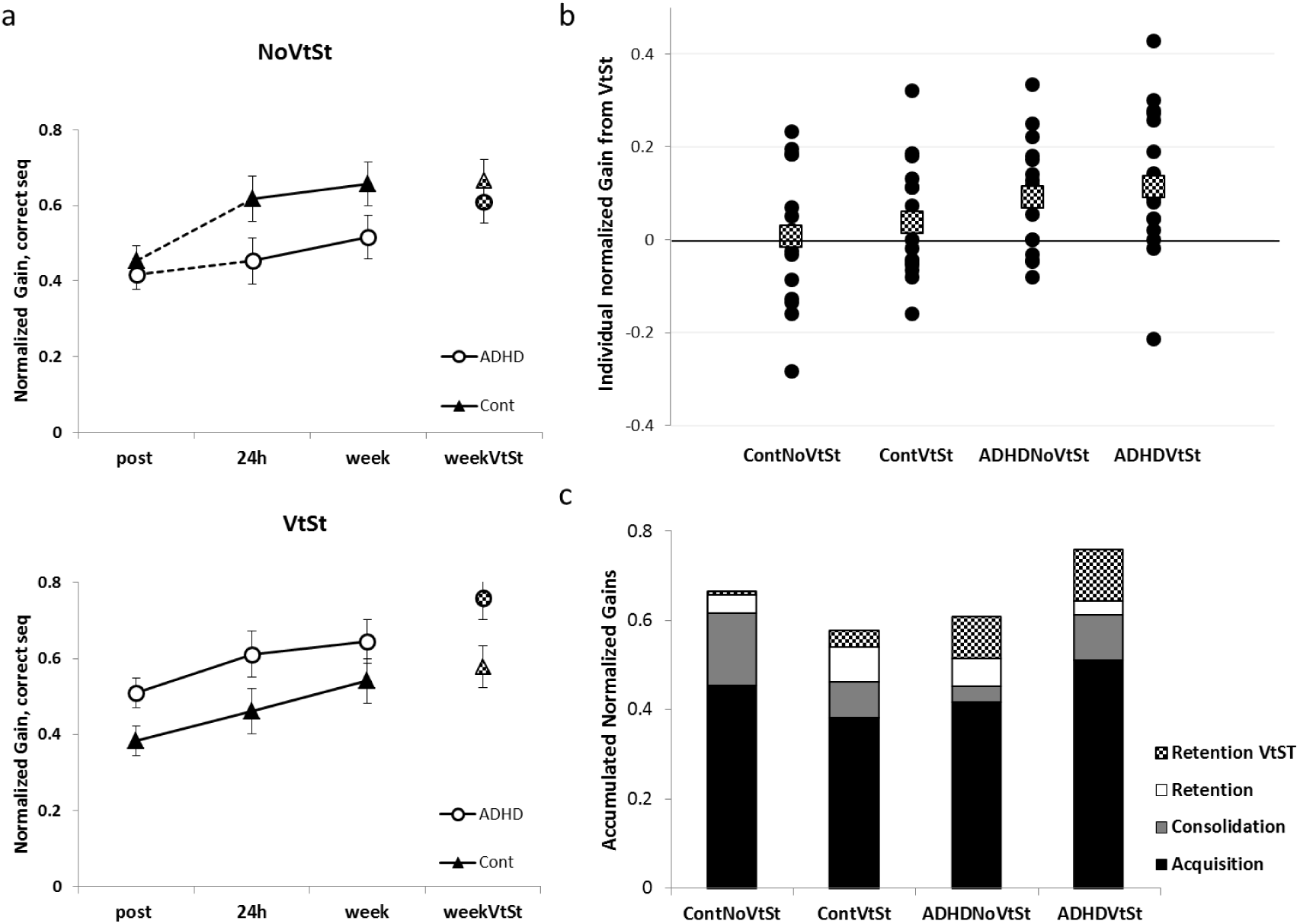
Normalized data. The gains at 3 time-points in participants trained with and without VtSt. (**a**) Group averages of normalized gains in performance speed, number of correct sequences (Δ) relative to each individual’s pre-training baseline performance. Each time-point represents the additional gains at the end of each of the 3 consecutive phases relative to baseline [post-test, acquisition=(post-pre)/pre; 24h, consolidation=(24h-pre)/pre; 1-week, retention=(retention-pre)/pre]. Lower panel - training with no VtSt; upper panel - training with VtSt). Dashed line - a significant interaction of group and time-point. Bars - SE. Circles - ADHD; triangles - Cont.; chess board markers - retention test with VtSt. (**b**) Individual normalized gains during the test with VtSt performed at 1-week retention (retentionVtSt). The individual VtSt gains scores were calculated as the difference between (the mean of the 4) test blocks with VtSt and the (4) test blocks without VtSt at retention; i.e., (retentionVtSt-retention)/pre-test. Squares chess-board markers - group mean relative gains. (**c**) Contribution of the three learning phases, and the affordance of VtSt during the final test, to the overall normalized gains in performance.

When training was afforded without VtSt there were, overall, significant gains in performance in terms of speed (number of correct sequences executed in the test) normalized to pre-test performance, at the 3 time-points (representing gains in performance at the end of 3 time-intervals: acquisition, consolidation, 1-week retention) in both groups (ContNoVtSt, ADHDNoVtSt) i.e., irrespective of whether participants had ADHD symptoms (F(2,60)=20.304, p<0.001, MSE=0.185, η^2^=0.404) **(Figure 4a, upper panel)**. There was no significant group effect (p=0.133), but there was a significant time-point X group interaction (F(2,60)=4.172, p=0.020, MSE=0.038, η^2^=0.122). Post-hoc comparisons indicated that the course of performance improvements differed significantly in the 24 hours post training consolidation phase; while both groups improved in terms of speed (F(1,30)=13.239, p=0.001, MSE=0.155, η^2^ =0.306), there was a significant interaction of group X time-points across the consolidation phase (F(1,30)=5.649, p=0.024, MSE=0.066, η^2^=0.158), originating from the ContNoVtSt group outperforming the ADHDNoVtSt group. Comparing the relative gains during the consolidation and retention phase in the 2 groups showed a significant main effect (F(1,30)=13.163, p=0.001, MSE=0.420, η^2^=0.305) and a trend towards a significant group effect (F(1,30)=3.838, p=0.059, MSE=0.378, η^2^=0.113), but no significant interaction (p=0.541). Thus, after the acquisition interval, if no VtSt was afforded, the Control and the ADHD groups had similar learning curves, yet the gap (differential gains) accrued consolidation interval was maintained **(Figure 4a, upper panel).**

Training with VtSt, however, resulted in a different pattern of results (**Figure 4a, lower panel**). Overall, significant gains in performance in terms of speed (number of correct sequences) normalized to pre-test performance, at the 3 time-points (representing the 3 time-intervals: acquisition, consolidation, 1-week retention) were observed for both groups (ADHDVtSt, ContVtSt), F(2,60)=20.276, p<0.001, MSE=0.179, η^2^ =0.395). There was no significant difference between the groups’ overall performance (p=0.135) and no interaction effects (p=0.491) were observed. However, a comparison of 2 time-points (post-test and 24 hours consolidation, representing the gains during the consolidation phase) showed that while both groups (ContVtSt, ADHDVtSt) had significant gains (F(1,31)=16.573, p<0.001, MSE=0.134, η^2^=0.348), there was a marginally significant group effect F(1,31)=3.414, p=0.074, MSE=0.315, η^2^=0.099), suggesting that the ADHDVtSt group was better than the ContVtSt group immediately after training and this advantage continued across the consolidation period. The group X time-point interaction was not significant (p=0.260). Both groups continued to improve over a 1 week interval. A comparison of 2 time-points (24 hours consolidation and 1-week retention) showed that the 2 groups (ContVtSt, ADHDVtSt) had significant gains (F(1,31)=6.299, p=0.018, MSE=0.052, η^2^ =0.169) during the retention phase but there was no significant group difference (p=0.174) and no significant interaction (p=0.312).

To directly compare the effects of adding the sensory stimulation during a performance test, in participants with and without ADHD, an rm-ANOVA with each participant’s normalized performance scores in the 2 test conditions (1-week, 1- weekVtSt) and the ADHD status (ADHD, Cont) was performed. There was a significant test condition effect (F(1,63)=11.877, p=0.001, MSE=0.117, η^2^=0.348) indicating overall better performance in the test blocks with VtSt, but also a significant interaction of test condition x ADHD status (F(1,63)=4.648, p=0.035, MSE=0.046, η^2^=0.070), indicating that participants with ADHD responded to the presence of the vibrotactile stimulation during the test in a different manner compared to non-ADHD controls. No group effect was found (p=0.697). Most of the participants with ADHD benefitted from the addition of VtSt during the testing of performance (**Figure 4b**). Post-hoc one-sample two-tailed t-test analyses showed that in both the ADHD groups, the mean additional gains in performance in the test blocks wherein background VtSt was afforded, were significantly above zero (t(16)=2.744, p=0.014; t(15)=2.695, p=0.017, ADHDVtSt and ADHDNoVtSt, respectively). However, in both Control groups the mean contribution of the added VtSt to performance was not significantly different from zero (t(15)=1.182, p=0.256; t(15)=0.213, p=0.834,ContVtSt and ContNoVtSt, respectively). Thus, the participants with ADHD also benefitted from the affordance of VtSt during performance testing, irrespective of whether they were exposed to the VtSt during training on the movement sequence a week earlier or not; no such group benefit was found in non-ADHD controls.

The groups’ mean normalized gains in performance accrued during specific time intervals along the course of learning the new motor sequence are presented in the **Figure 4c**. Following the single training session, all groups improved by more than 50% relative to the pre-training performance baseline. But by the end of the study (as expressed in the test session at 1 week post-training) whether participants were given training with or without VtSt had a differential effect on the overall gains in performance, depending on whether the trainees had ADHD symptoms or not. After training with no VtSt, participants with ADHD showed an overall improvement of performance speed (by 51.6%) but the gains were, on average, smaller than those attained by their typical peers with no ADHD (overall improvement, 65.8%) who trained in the same condition. However, VtSt during training (in an otherwise identical protocol) benefited subsequent performance in participants with ADHD (overall improvement, 64.5%) but was relatively detrimental for typical controls (overall improvement, 54.1%) (**Figure 4c**).

A performance advantage for the participants with ADHD symptoms was also apparent, irrespective of how they were trained, when performance was tested in the presence of VtSt (during the retention test). VtSt during the test boosted the performance in the majority of the participants with ADHD (mean additional gain of 11.4% and 9.2% in ADHDVtSt and ADHDNoVtSt, respectively), while in typical controls the mean effect was not significant (mean additional gain of 3.7% and 0.7% in ContVtSt and ContNoVtSt, respectively). Note, however, that in both groups, there were individuals that responded to the presence of the VtSt, either by boosting or by degrading their speed of motor sequence performance (**Figure 4b**).

### Chronotype and Sleep data

There was a trend towards a significant difference in the mean MEQ scores between the ADHD and the control participants (**Table 1**), with persons with ADHD more inclined to be evening-oriented. This tendency was observed in spite of the fact that extreme morning and evening chronotypes were excluded from the experiment. Means of time-in-bed, sleep latency (time to fall asleep), total sleep time (minutes), and sleep efficiency ((total sleep time/time in bed)*100) parameters were derived from the actigraphy during the post-training night. These parameters were compared across participants with and without ADHD using two-tailed independent sample t-tests. No significant differences were found between the ADHD and control participants with the exception of sleep latency; participants with ADHD had a marginally significant tendency to have longer sleep latencies (**Table 1**).

**Table 1.** MEQ scores and Actigraphy data.

|  | ADHD / Cont<br>(mean±SD) | t (62) | Sig. (2-tailed) |
| --- | --- | --- | --- |
| **MEQ score** | 48.4±7.6 / 51.8±6.9 | -1.846 | **0.070** |
| **Sleep Latency, min** | 18.9±25.6 / 9.6±14.4 | 1.697 | **0.095** |
| **Sleep Efficiency, %** | 75.6 / 77.2 | -0.466 | 0.643 |
| **Time in bed, min** | 480.1±91.2 / 456.6±75.0 | 7.075 | 0.285 |
| **Total Sleep Time, min** | 358.2±72.7 / 361.7±66.2 | -0.191 | 0.849 |
| **Mean Number of Awakenings** | 5.3±3.6 / 4.8±2.5 | 0.959 | 0.341 |

Pearson correlation analyses showed that in the NoVtSt groups, irrespective of the ADHD status, higher MEQ scores (higher morningness) correlated with higher consolidation gains (r=0.34, n=33, p=0.032). In the groups afforded the VtSt no such correlation was found (r=-0.25, n=33, p=0.161).

## Discussion

In typical young adults minor task-irrelevant vibro-tactile stimulation afforded during training on a novel sequence of movements can selectively impair off-line consolidation processes and, as a result, decrease the long-term practice related gains in performance^29^. Here we replicated this result, but, we also show that in the same background sensory stimulation condition there was, paradoxically, a positive effect on learning and the generation of long-term practice related gains, in young adults with ADHD. Moreover, the current results demonstrate an experiential condition wherein persons with ADHD have a clear advantage in skill acquisition and in subsequent performance over their typical peers.

In line with previous studies^1,10,30^, participants with ADHD, who had practice without sensory stimulation (NoVtSt), i.e., when training in standard, quiet, laboratory conditions, showed, in comparison to their typical peers (without ADHD), less-than-expected overnight consolidation-phase gains in the performance of the trained movement sequence. Participants with ADHD tended to underperform when tested overnight and began to lag behind typical peers at later time-points, despite the fact that both groups expressed equal within-session gains. However, the current results show that an identical practice protocol, but with VtSt afforded in the background throughout training, resulted in larger improvements in task performance within-session, as well as in robust overnight consolidation phase gains in persons with ADHD. These gains were well retained and expressed in a re-test after a week-long interval in which no additional training was afforded. Typical young adults, training with VtSt, showed clear costs in terms of their ability to express overnight delayed gains in performance as well as by the end of the study period. Moreover, by the end of the study period, when performance was tested with VtSt afforded during the tests, participants with ADHD were clearly helped; many of their typical peers were hampered. Thus, training with background vibratory noise resulted in distinct and opposing effects in young adults with and without ADHD; while the learning related gains in performance tended to diminish in typical adults, participants with ADHD became better learners, expressed larger consolidation phase gains and subsequently outperformed their typical peers. The affordance of VtSt during the subsequent testing of performance benefitted participants with ADHD but not their typical peers.

The addition of vibrotactile-auditory sensory stimulation (VtSt) had no effect on the immediate post-training performance of the task by participants without ADHD. The absence of adverse effects on performance and on the magnitude of ‘online’ learning in typical participants suggests that the background stimulation was indeed experienced as minimal and did not significantly avert attention from the task during the training session. Participants in the ADHD group also showed no sign of being distracted by the background stimulation; in fact the VtSt stimulation afforded during practice turned the practice session into a more efficient learning experience for them.

Eveningness and sleep problems are common in adults with ADHD^31^. However, the actigraphy data showed that in the current study the participants of the ADHD and the Cont groups had, overall, similar sleep profiles in the post-training night, except for the tendency of the ADHD participants to have longer sleep latency periods. The similarity between groups in terms of sleep parameters was partly the result of the screening procedure adopted, because extreme morning and evening chronotypes were excluded from the experiment. As no differences were found between the typical controls and the participants with ADHD, our results cannot be taken to reflect a bias in sleep parameters. Note that all participants had relatively low sleep efficiency (percentage of time in sleep relative to total time in bed). This is an increasingly common finding in young adults and adolescents, attributed to the use of light-emitting devices at evening hours^54^. We also tested for a possible relationship between the participants’ chronotype and their learning abilities, given that training and testing took place in the morning hours^31^. Evening oriented participants are more likely to show lower arousal levels and lower cognitive performance^55^ during the morning hours compared to morning-oriented peers, and we conjectured that this may lead to a less engaging learning experience and subsequently to the expression of smaller delayed gains in performance. Indeed, the results showed that in the NoVtSt groups, higher morningness correlated with larger consolidation phase gains irrespective of whether participants had ADHD. Because of the differential effects of VtSt in the 2 groups and the small number of individuals in each of the 2 VtSt groups, correlation analyses with chronotype in these samples were not informative.

The impact of ‘state’^33^ (ongoing or background activity^56^), and, specifically, levels of arousal, prior to or during test or learning sessions is, in practice, an often neglected factor in memory research. Even in standard laboratory training protocols an optimal arousal state cannot be, but often is, assumed^10,33,57^. Moreover, individuals may differ in the level of arousal optimal for enabling them to attain optimal performance^33^ and, as the Yerkes–Dodson model suggests, both too little or too much arousal can adversely affect task performance^38^. Vibratory stimulation is considered an alerting intervention, improving vigilance^58^ and increasing skeletal muscle tone. In addition, vibratory or auditory^59^ stimulation can induce affective reactions^60^. Both the enhancement or impairment of memory^25^ have been shown to be modulated by stress^61,62^ depending on task and training conditions. Specific combination of hippocampal activation during motor sequence practice session and of post-training night sleep may be a pre-requisite for promoting the expression of delayed gains in motor sequence task performance during the consolidation phase^63^. Thus, both background conditions and the individuals ‘state’ during and after the performance of a given task may affect (as “gating” factors) the acquisition and, importantly, the consolidation of skills (‘how to’ knowledge)^10,29^.

In ADHD arousal regulation may be atypical and thus may constitute one of the ‘core’ characteristics of the condition^64^. Individuals with ADHD tend to be under-aroused^39,42,43^, and often experience difficulty in sustaining attention during repetitive tasks^51^. The restless behaviour of individuals with ADHD has been interpreted as self-stimulation in order to raise their arousal level^50^ and, consequently, performance.

In healthy adults vibratory stimulation was reported to neutrally or negatively affect attention and cognition^65,66^. In clinical populations, as in ADHD, background stimulation, vibration or white auditory noise^50^ have been proposed as means to enhance attention, and benefit learning processes^67^, and were even suggested as an adjunct in enhancing motor training^68^ and rehabilitation^69^. The current results are in line with and extend these notions. We propose that the observed benefits of VtSt to participants with ADHD may relate to upregulated arousal due to the concurrent sensory stimulation. Note, however, that in the presence of VtSt there were individuals that benefited from stimulation, also among control participants; other individuals were severely interfered by it. Thus, the individual’s arousal state, as well as sensory responsivity to vibratory stimulation, may be predisposing factors in determining whether one would benefit or lose from the presence of background-environmental noise.

There is evidence that non-pharmacological interventions may up-regulate skill learning in ADHD, with recent studies focusing specifically on motor learning. First, some of the relative learning deficits in persons with ADHD could be corrected when training was shortened^1,2,6,3^ presumably by decreasing the burden of long repetitive practice on mechanisms of sustained attention^8^. Second, motor training scheduled to evening hours was found to enhance off-line memory consolidation in young adults with ADHD and the learning gap vis-à-vis typically developing adults was closed ^10^. Evening hours are more suitable in terms of arousal levels for evening-type individuals, such as many of the individuals with ADHD^31^. Related to this notion is the finding that five minutes of vigorous physical activity can improve affect and xecutive functioning of children with symptoms of ADHD^32^.

To conclude, our results suggest that: i) procedural memory acquisition and consolidation processes are extant in young adults with ADHD and this potential can be best unveiled in specific bio-behavioural conditions; ii) such bio-behavioural conditions should be afforded during training to enhance learning in ADHD; iii) minor background vibro-tactile stimulation may constitute an effective aid during procedural learning in ADHD; in typical peers it may slow or dampen consolidation processes. The current results also underscore the possibility that even temporary failures of arousal in ADHD can result in long-lasting and accumulating deleterious effects. We conjecture that many behavioural difficulties expressed in individuals with ADHD are related to under-arousal and that these deficits can be compensated by manipulating physical conditions so as to increase levels of arousal. From a different perspective, our results suggest that ADHD can be considered a neuro-behavioural phenotype that may confer advantages in performance, learning and skill memory consolidation in ‘noisy’ conditions that adversely affect typical non-ADHD peers.

## Methods

The study was approved by the Human Experimentation Ethics committee of the University of Haifa. All participants signed an informed consent form in accordance with the Declaration of Helsinki prior to the start of the experiment. Participants were paid (150 NIS, approximately $40) for their participation.

Sixty five right-handed young participants (aged 19-35, 24.6±3.8 mean ± s.d., 24% males) enrolled in the study. The sample size was based on the effects observed in previous studies with typical^15,29^ and ADHD^10,30^ participants, where the groups that did not experience interfering interventions showed 20-30% gains in performance speed during the consolidation phase, whereas the groups in suboptimal training conditions showed less than 7% delayed gains. As we expected the ADHDNoVtSt and the ContVtSt groups to have deficient off-line learning, and the ADHDVtSt and the ContNoVtSt groups to have normal off-line learning, we expected to observe similar differences in delayed gains in performance. Based on a STD of 15% in each group, and a power of 0.80, we required 16 subjects per group^70^.

Participants were recruited through the University of Haifa and the Technion’s mass media platforms (University newspaper, Facebook pages) and an electronic message sent through the university’s Centre for Students with Disabilities, for a “study on motor learning and memory”. Thirty three participants met the inclusion criteria for ADHD group, and thirty two typically developing adults matched by age and education, served as a control group (Controls). Inclusion criteria for the ADHD groups were as follows: (1) a formal psycho-didactic diagnosis of an attention deficit disorder (either ADD or ADHD) from an authorized clinician, psychiatrist or neurologist, approved by the University Centre for Students with Disabilities within 5 years of the current study; (2) a positive screening on the adult ADHD self-report scale (ASRS)^71^ and (3) no stimulant treatment for ADHD (methylphenidate or other stimulant drugs) during the recent period (>month). The participants of the ADHD group responded positively on 11 out of 18 items of the ASRS on average (10.9±2.8, mean±s.d.). The control participants met ≤ 3 out of 6 criteria of the ASRS screening questionnaire (first 6 items). All control participants affirmed that they were not suspected (by family members or teachers) to have, and were never diagnosed as having, ADHD/ADD during their childhood or adulthood.

Pre-screening was done by a short telephone interview (including questions of general health status and basic demographic data) to exclude persons with diagnosed sleep, neurological or psychiatric disorders (other than ADHD), motor-skeletal diseases, use of drugs, heavy alcohol consumption and regular smoking; also excluded were persons reporting “blind” typing or skilled musical instrument playing and those reporting less than 6 hours of sleep per night or defining their sleep quality as low and insufficient. Prior to the commencement of the experiments, the invited participants completed the PSQI sleep questionnaire; only participants with global scores below the cut-off (≤5) were included^72^. All participants underwent chronotype assessment using the Horne-Östberg Morningness-Eveningness Questionnaire (MEQ)^73^ extreme chronotypes (scores >70 – extreme morningness and <30 – extreme eveningness) that could bias motor performance, were excluded.

The participants were trained and were tested on an explicitly instructed five-element finger-to-thumb opposition sequence (FOS), as in^29^. All sessions took place during morning hours, between 8:00 and 12:00 (Figure 1a). The participants were seated with their task performing arm positioned on a table, comfortably extended, with the palm facing up to allow video recording of all finger movements. Visual feedback was not allowed; the participants were instructed to avert their gaze away from the performing hand. Headphones (Beats Pro) were used throughout the experiment.

The experiment included three sessions; the general design of the study is schematized in Figure 1b. In the first session, the experimenter explained and demonstrated the thumb-to-finger opposition movements the sequence assigned to each individual for training (Sequence A, Figure 1a). The participant had to correctly perform the instructed sequence on three consecutive self-paced iterations (warm-up and as a check to ensure knowledge of the required movement sequence), otherwise the instructions and demonstration were repeated, and then underwent the pre-training performance test (pre-test). The pre-test was followed by a cued, structured, training session, and an immediate post-training test (post-test). Each performance test consisted of four 30 sec long blocks. Before each block the participant was reminded to perform the sequence repeatedly and continuously “as fast and as accurately as possible” during the interval denoted by a start and a stop auditory cue signals, delivered through headset. Participants were instructed that if they became aware of committing an error they should immediately continue to the required sequence. Each test block was followed by a 30 sec rest interval. The training session consisted of 160 cued repetitions of the sequence, afforded in ten 30 sec. blocks. Thirty sec. long rest intervals were afforded between blocks. In each of the training blocks, the initiation of each sequence repetition was cued by a brief sound delivered at a rate 2.5 sec. per sequence.

After completing the training blocks, participants were asked to perform four additional test blocks (post-test), with instructions identical to those provided before the pre-test. Participants were re-tested, each re-test consisting of four continuous performance blocks at 24h, following a night sleep (24h re-test) and at one week (week re-test) following the training session. In addition, at the week re-test, all participants were tested in four continuous performance test blocks with the vibratory stimulation afforded concurrently, to test the on-line effect of sensory stimulation.

In pilot experiments, a higher level of stimulus intensity (~65dB) was judged as distracting by some of participants with no ADHD. Four participants with no ADHD (not included in the main study) were interviewed as to the discomfort induced by vibration stimulation at ~41dB. The level of vibratory stimulation was judged as minimally uncomfortable and non-distracting in two conditions, stimulation provided with and without the performance of the motor task. In addition, the recorded sound resulting from the cushion’s vibrations was played back through the headphones at 40dB. The participants in the NoVtSt condition were seated on the same cushion with the current switched off. All auditory signals, the auditory cues for the initiation and termination of each test and training block and the continuous auditory background vibration sounded during the training blocks in the VtSt groups, were recorded and provided using Audacity program (Ver 2.2, GNU General Public License).

Participants were instructed to concentrate on the motor training task to maintain maximum accuracy in sequence execution irrespective of the presence of background stimulation. At the end of each session participants were instructed not to repeat or practice the movement sequence they were trained on between the meetings.

Participants wore an actiwatch (Actigraph Co.) for 24 hours, starting from the end of the post-test to monitor sleep time, quality and length during the post-training night. The data were analysed using the ActiLife 6 software.

Performance data were analysed from video recordings. Measures of *speed* (number of correct sequences) and *accuracy* (number of errors) of performance at each 30-sec test block were derived. Means of the performance in the 4 test-blocks at each of the 4 time-points (pre-test; post-test, 24h test, week retention test) as well as in the test at 1-week post-training with VtSt afforded, were calculated. In addition, normalized data (relative improvement) for speed after the acquisition, the consolidation and the retention intervals were calculated relative to the mean pre-test, baseline, performance of each individual. Absolute and normalized speed and accuracy performance scores were analysed separately. Independent samples, 2-tailed t-tests were used to compare between the pre-test performance levels of the groups. Repeated measures analysis of variance (rm-ANOVA) with the 4 time-points as a within-subject factor and group (ADHDNoVtSt, ADHDVtSt, ContNoVtSt, ContVtSt) as a between-subjects factor were conducted to assess the changes in performance across the study period. Post-hoc rm-ANOVAs comparing pairs of consecutive time points were conducted to test performance changes across specific phases: acquisition (pre-test vs. post-test), consolidation (post-test vs. 24h test) and retention (24h test vs. week). The affordance of VtSt during the performance test was assessed using rm-ANOVAs with 2 test conditions (with and without background VtSt) as a within-subject factor and group (ContNoVtSt, ContVtSt) as a between subjects factor.

## Author contributions

MK, LM and AK conceived and designed the experiments. LM and NA collected the data. LM analyzed the raw data. MK and LM did the statistical analyses and MK, LM and AK were responsible for the interpretation of the data. MK, LM and AK wrote the article.

